# A bivalent CMV vaccine formulated with human compatible TLR9 agonist CpG1018 elicits potent cellular and humoral immunity in HLA expressing mice

**DOI:** 10.1101/2022.03.02.482616

**Authors:** Vijayendra Dasari, Kirrilee Beckett, Shane Horsefield, George Ambalathingal, Rajiv Khanna

## Abstract

There is now convincing evidence that the successful development of an effective CMV vaccine will require improved formulation and adjuvant selection that is capable of inducing both humoral and cellular immune responses. Here, we have designed a novel bivalent subunit vaccine formulation based on CMV-encoded oligomeric glycoprotein B (gB) and polyepitope protein in combination with human compatible TLR9 agonist CpG1018. The polyepitope protein includes multiple minimal HLA class I-restricted CD8^+^ T cell epitopes from different antigens of CMV. This subunit vaccine generated durable anti-viral antibodies, CMV-specific CD4^+^ and CD8^+^ T cell responses in multiple HLA expressing mice. Antibody responses included broad T_H_1 isotypes (IgG2a, IgG2b and IgG3) and potently neutralized CMV infection in fibroblasts and epithelial cells. Furthermore, polyfunctional antigen-specific T cell immunity and antiviral antibody responses showed long-term memory maintenance. These observations argue that this novel vaccine strategy, if applied to humans, could facilitate the generation of robust humoral and cellular immune responses which may be more effective in preventing CMV-associated complications in various clinical settings.

## Introduction

Human cytomegalovirus (CMV) is a betaherpes virus and approximately 60% of adults in developed countries and more than 90% in developing nations are infected with CMV (1). Primary CMV infection in healthy individuals is usually asymptomatic or clinically manifested with mild symptoms. However, individuals with compromised immune system, such as transplant patients and people infected with HIV, are at increased risk of developing CMV-associated complications. During pregnancy, depending on the serostatus, CMV can transmit vertically from mother to foetus as a result of primary infection, superinfection or reactivation of latent virus and cause congenital infection, neurodevelopmental delay, microcephaly or neonatal hearing loss in childhood (2, 3).

CMV has high cell tropism and enters by fusing its envelope glycoproteins, such as gB, gH, gL, gO and UL128/UL130/UL131A with either plasma membrane or endosomal membrane (4, 5). The gB protein is a highly conserved envelope protein, during CMV infection it acts as a fusogenic molecule in association with the gH/gL/gO protein complex. In healthy individuals 70% of total CMV-specific serum antibodies are directed against gB protein and neutralising antibodies against gB protein was shown to inhibit cell-to-cell virus spread and entry of cell-free virus in fibroblasts, endothelial and epithelial cell types (6, 7). CMV infection in healthy individuals induces strong and diversified cellular immune responses which includes high frequencies of CMV-specific CD8^+^, CD4^+^ and γδ T cells (8). Lack of CMV-specific cellular immunity in transplant patients has been shown to have an augmented occurrence of CMV replication episodes and CMV-associated complications. However, over the years adoptive transfer of CMV-specific CD4^+^ and CD8^+^ T-cells has been shown to have a profound benefit in controlling CMV-associated disease in hematopoietic stem cell transplant (HSCT) and solid organ transplant (SOT) recipients (9-14). In addition, compromised cellular immunity can impair immune control of CMV infection in children with congenital infection and postnatal CMV infection (9). Collectively, these observations suggest that CMV-specific humoral and cellular responses are key components of protective immunity and an effective CMV vaccine would need to induce both humoral and cellular immunity against multiple antigens. Indeed, a wide array of CMV vaccines have been developed using a number of delivery platforms, such as live attenuated, subunit, DNA plasmid, mRNA and recombinant viral vectors and tested in a number of preclinical and clinical trials (15). However, successful licensure of a CMV vaccine formulation remains elusive.

In this study, we have engineered a novel artificial recombinant polyepitope protein (CMVpoly), which includes 20 HLA class I-restricted T cell epitopes from multiple CMV antigens expressed as a “string of beads”. We have also engineered CMV gB protein and developed an efficient protein production and purification process to obtain gB as a multimer in solution. CMVpoly and gB protein in combination with human compatible adjuvant CpG1018 was formulated as a bivalent CMV vaccine with the aim to induce CMV-specific cellular and humoral immunity. Our results demonstrated that immunisation with CMV vaccine consistently generated robust CMV-specific neutralising antibodies, CD4^+^ and CD8^+^ T cell responses. More importantly, long-term follow-up analysis showed that the CMV vaccine can induce durable CMV-specific humoral and cellular immune responses.

## Results

### CMV antigen design

To target a broad repertoire of immunodominant CMV antigens we designed a vaccine formulation that included recombinant CMVpoly and CMV-gB proteins in combination with human compatible adjuvant, CpG1018. The CMVpoly protein is designed to encode 20 HLA class I restricted cytotoxic CD8^+^ T-cell epitopes from five highly conserved immunodominant antigens (pp65, IE-1, pp150, pp50 and DNAse) (Table 1). CMVpoly targets 16 different HLA class I alleles and they cover 94% of a multi-ethnic population worldwide. To optimise the processing and presentation of CMVpoly protein through HLA class I pathway proteasome liberation amino acids (AD or K or R) were added at the carboxyl-terminus of each epitope (Table 1) (16). CMVpoly protein was expressed using *E. coli* BL21-codonPlus (DE3) RP protein expression host and was purified using SP-Sepharose and Q-Sepharose chromatography columns (Fig. 1a). To assess processing and presentation of T cell epitopes within CMVpoly, peripheral blood mononuclear cells (PBMCs) from 10 healthy virus carriers were incubated with CMVpoly protein and then cultured for 14 days in the presence of recombinant IL-2. All 10 healthy virus carriers showed expansion of multiple CMV epitope-specific T cells (Fig. 1b) and these T cells showed polyfunctional profile and cytotoxicity against CMV-infected cells (Fig. 1c & d).

**Table 1:**
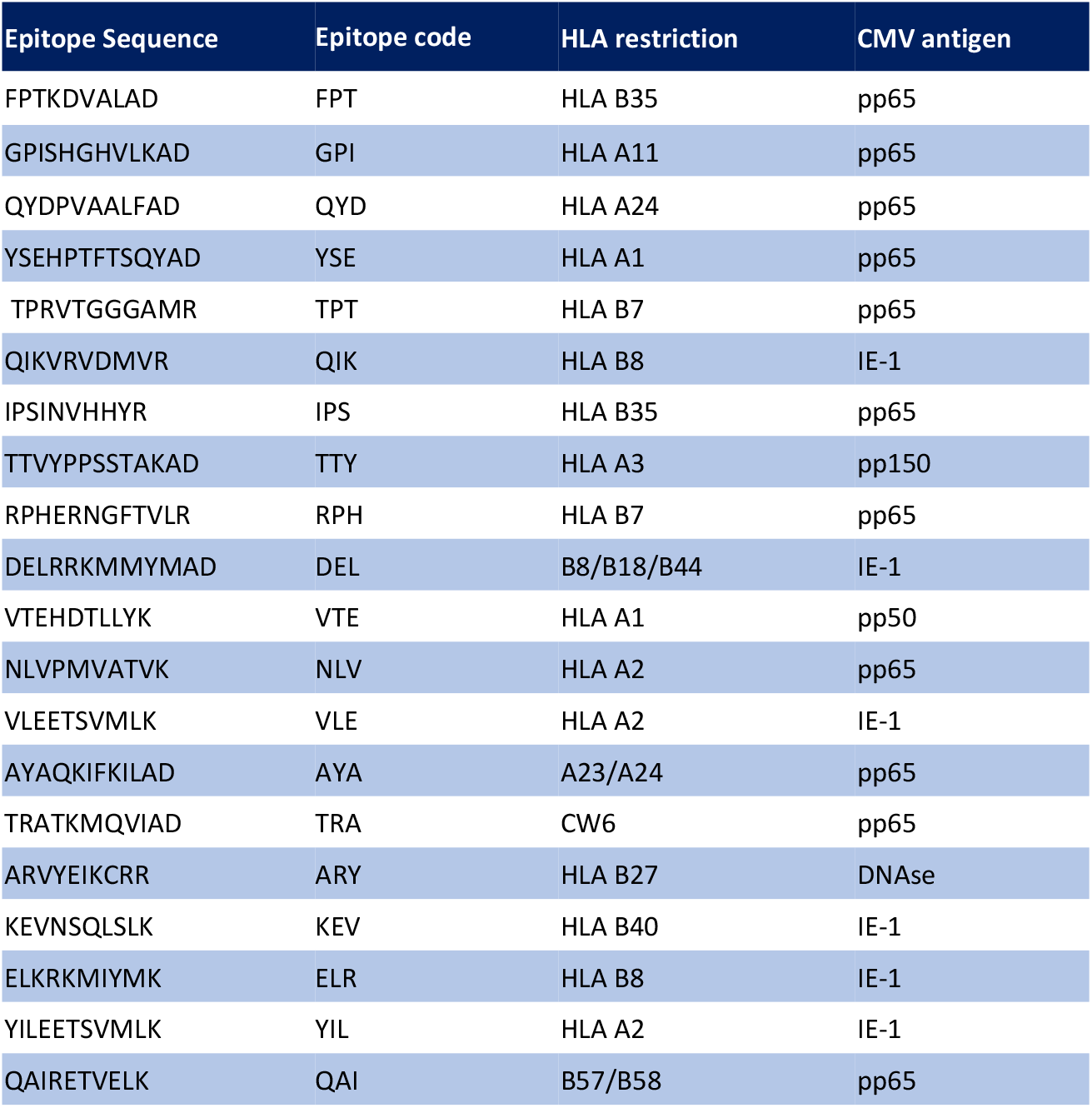
List of HLA class I-restricted CD8^+^ T cell epitopes included in the CMVpoly protein

**Fig 1.**
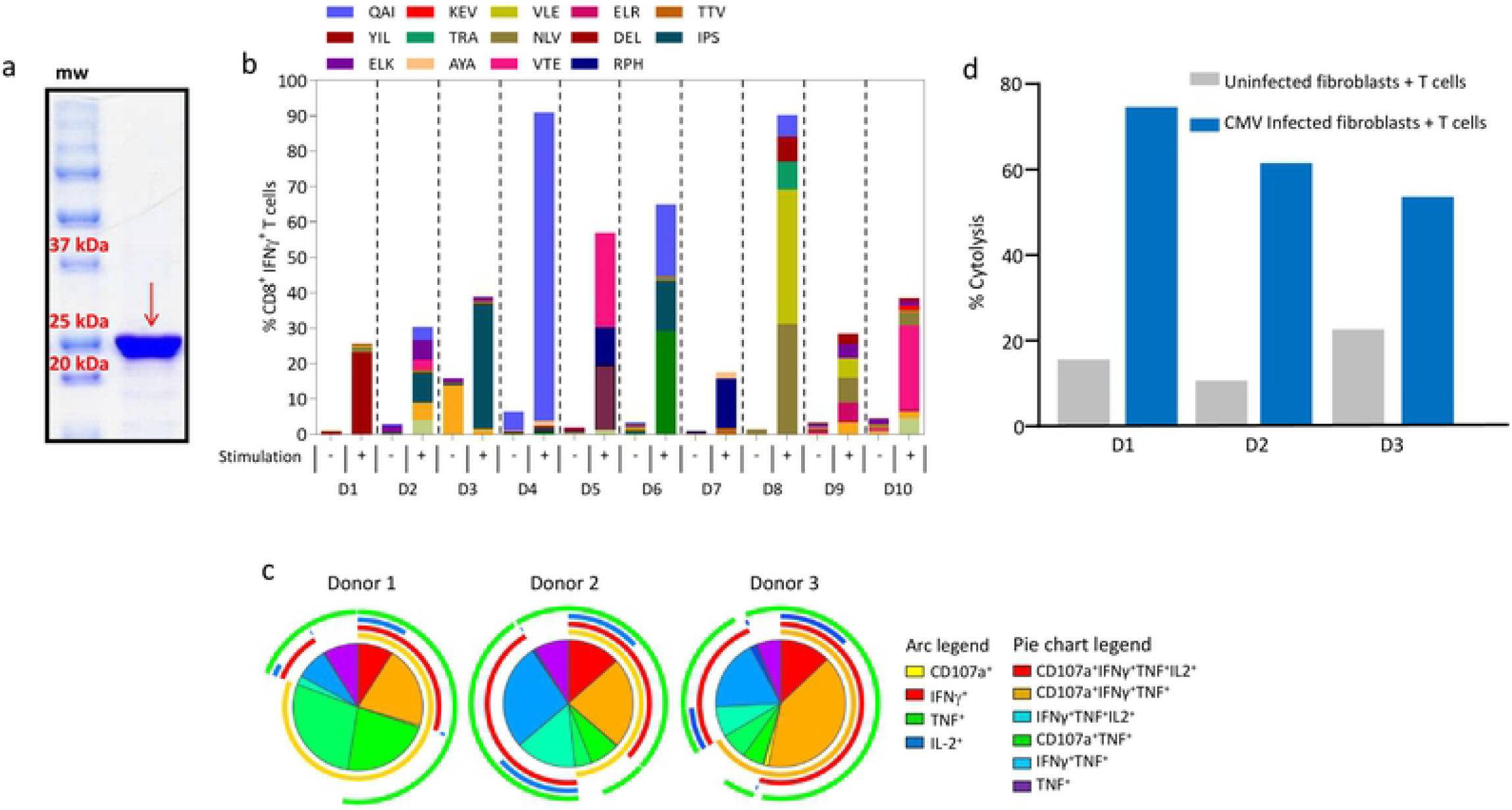
CMVpoly protein purification, characterisation and *in vitro* immunogenicity analysis. (a) SDS-PAGE showing purified CMVpoly protein with a molecular weight of 25.1 kDa. (b) *In vitro* expansion of CMV-specific CD8^+^ T cells from healthy CMV seropositive donor PBMCs following stimulation with CMVpoly protein. Briefly, PBMCs from healthy donors were stimulated with CMVpoly protein (20μg) and cultured for 14 days in the presence of IL-2. These *in vitro* expanded T cells were assessed for CMV epitope-specific reactivity in an intracellular cytokine assay. Bar graph represent the comparative frequencies of IFN-γ expressing CMV-specific CD8^+^ T cells in unstimulated PBMCs and *in vitro* expanded T cells. (c) Pie chart shows polyfunctional profile of CMVpoly stimulated CD8^+^ T cells comprised of various combinations of CD107a^+^, IFNγ^+^, TNF^+^ and/or IL-2^+^. Arc indicates the proportion of CMV-specific CD8^+^ T cells producing individual function: CD107a (yellow) IFNγ (red), TNF (green) and IL-2 (blue). (d) Antigen-specific CD8^+^ T cells expanded with CMVpoly protein kill CMV-infected fibroblasts. Human fibroblasts expressing HLA A*01:01, A*02:01 and B*08:02 were infected with CMV TB40/E strain and then exposed to CMVpoly stimulated CD8^+^ T cells from three different donors at an effector to target ratio of 1:1 and for 48 hours and the cytotoxicity response was monitored in real time using xCELLigence and percent lysis determined. Uninfected fibroblasts were used as control in this assay.

CMV gB amino acid sequence from AD169 strain was designed to express extracellular domain (aa 24 to 700) and intracellular domain (aa 776 to 906) as a contiguous sequence. The membrane proximal and transmembrane regions were deleted and native furin cleavage site at aa 456-459 was mutated (RTRR to QTTQ) to enhance the recombinant gB protein secretion (Fig 2a). The CMV gB was expressed in CHO cells and then purified using anion, CHT type II and cation exchange chromatography techniques and purified protein was further characterised by size exclusion chromatography and SDS-PAGE analysis (Fig. 2b). Mass photometery analysis showed that purified CMV gB protein existed as oligomers in solution comprising two species with molecular masses of 398 kDa and 796 kDa, indicating the composition was predominantly a trimeric form of gB in addition to dimers of trimeric gB (Fig. 2c). Oligomeric gB protein was also observed using negative-stain electron microscopy (Fig. 2d).

**Fig 2:**
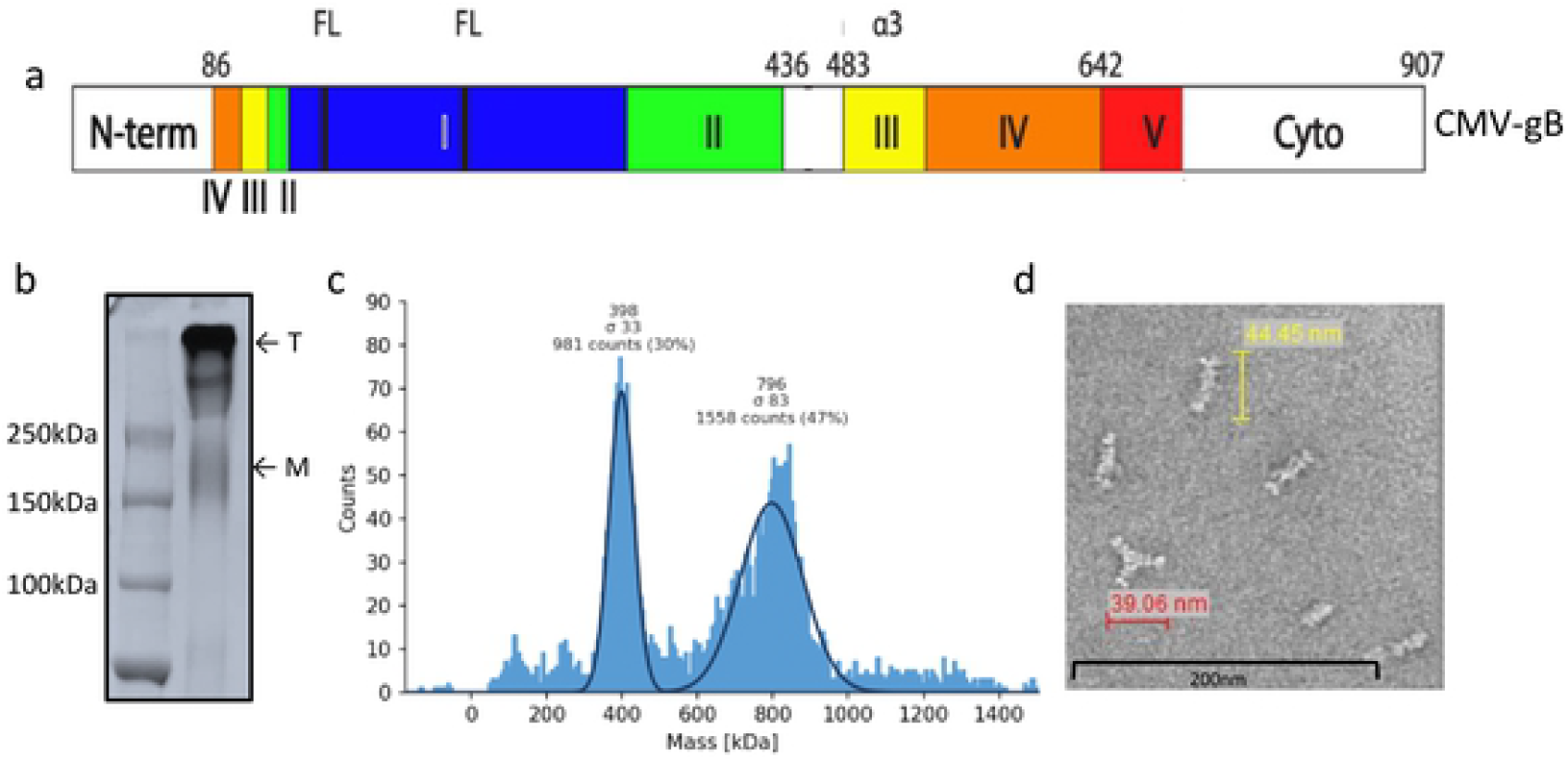
CMV gB protein purification and characterization: (a) Schematic representation of CMV gB protein domains included in the expression vector for recombinant protein expression. (b) SDS-PAGE analysis of purified CMV gB protein under non-reducing condition. Protein migrates as a trimer or dimer of trimer (T). A small fraction of monomer (M) is also visible on SDS-PAGE. (c)Mass photometry analysis of CMV gB protein confirming trimer and dimer of trimer form of gB protein; molecular weight for trimeric gB was 398 kDa and 796 KDa for dimer of trimer. (d) Negative stain electron micrographs of CMV gB protein representing oligomeric forms of gB protein. Scale bar, 200 nm.

### Assessing CMV vaccine immunogenicity in multiple HLA expressing mice

CMVpoly and gB proteins were formulated with human compatible CpG1018 adjuvant, a type B oligodeoxynucleotides, which acts as a toll-like receptor 9 agonist and has been shown to enhance immunogenicity in various preclinical and clinical trials and also approved for human use (17, 18). In the first set of experiments we assessed the CMV vaccine immunogenicity in multiple HLA expressing mice (HLA A1, A2, A24, B8 and B35). These mice were immunised subcutaneously with two dose prime-boost regimen (day 1 and 21), spleens and blood samples were collected seven days after the booster dose to assess CMV-specific humoral and CD4^+^ and CD8^+^ T cell responses (Fig. 3a). CD8^+^ T cell responses were recalled with CMV peptide epitopes restricted through HLA A1 (VTE and YSE), A2 (NLV, VLE and YIL), A24 (QYD and AYA), B8 (QIK, ELR and ELK) and B35 (FPT and IPS) alleles. CMV-specific CD4^+^ T cell responses were recalled with an overlapping gB pepmix. *Ex vivo* analysis of cellular immune responses indicated that most of the HLA expressing mice generated a broad range (0-2.5%) of antigen-specific CD8^+^ and CD4^+^ T cell responses (Fig. 3b & c). *In vitro* stimulation of splenocytes with HLA matched CMV CD8^+^ T cell epitopes induced 40 to 200 fold expansion of antigen-specific CD8^+^ T cells from all HLA expressing mice (Fig. 3d). These expansions were significantly higher in HLA A1, HLA A24 and HLA B8 expressing mice when compared to HLA A2 and B35 expressing mice (Fig. 3d). *In vitro* stimulation of splenocytes with gB overlapping peptides also triggered robust expansion of gB-specific IFNγ producing CD4^+^ T cells in all HLA expressing mice when compared to placebo control mice. Significantly higher frequencies of gB-specific CD4^+^ T responses were observed in HLA A24 mice when compared to HLA A1, HLA A2, HLA B8 and HLA B35 expressing mice (Fig. 3e). No significant differences in gB-specific CD4^+^ T responses were observed between mice expressing HLA A1, A2, B8 and B35 (Fig. 3e). Previous studies have shown that protective immunity mediated through CD4^+^ and CD8^+^ T cells is associated with production of multiple effector cytokines and impaired polyfunctionality of CMV-specific T cells can reduce control of CMV replication (19-21). Polyfunctional analysis revealed that the majority of CMV-specific CD8^+^ and CD4^+^ T cells from HLA expressing mice expressed multiple cytokines including IFNγ, TNF and IL2 or IFN-γ and TNF (Fig. 4).

**Fig 3:**
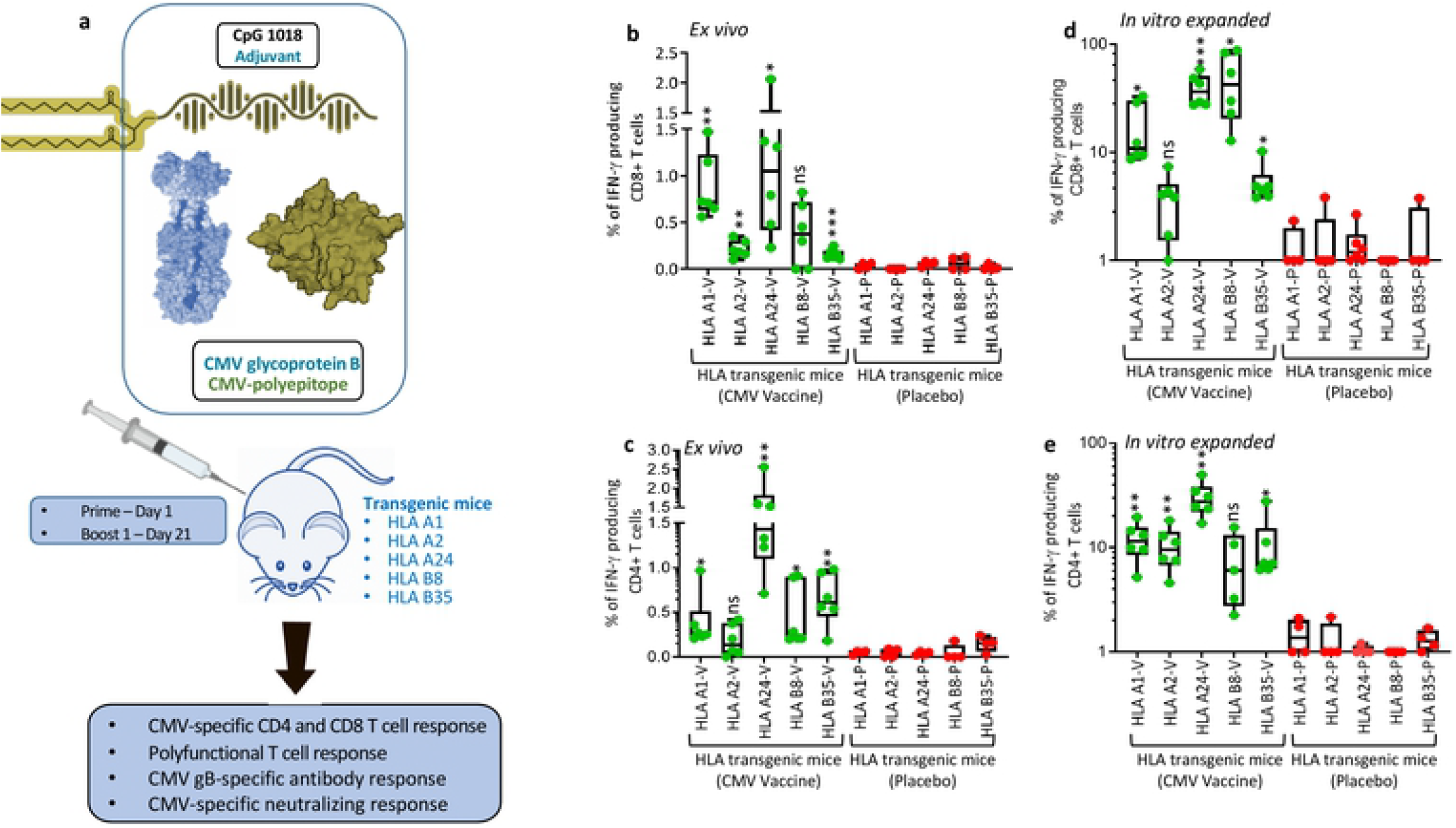
CMV vaccine immunogenicity evaluation in multiple HLA expressing mice: (a) Schematic outline of CMV vaccine immunogenicity evaluation in HLA expressing mice. CMV vaccine formulation included CMVpoly (30µG) and gB (5µg) proteins admixed with CpG1018 (50µg) adjuvant. Five different human HLA expressing mice (HLA A1, A2, A24, B8 and B35) were immunised subcutaneously with CMV vaccine (N = 6) or CpG1018 alone (N = 4) as a placebo control on days 0 and 21 and CMV-specific cellular and humoral immune responses were assessed on day 28. (a and b) *Ex vivo* CMV-specific CD8^+^ and CD4^+^ T cell responses in HLA A1, A2, A24, B8 and B35 expressing mice immunized with CMV vaccine. Splenocytes from immunized mice were stimulated with HLA matched CD8^+^ T cell epitopes (Panel a) or CMV gB overlapping pepmix (Panel b) and then assessed for IFN*γ* expression using intracellular cytokine assays. (d and e) *In vitro* expansion of CMV-specific CD8^+^ and CD4^+^ T cells from splenocytes of immunized mice following *in vitro* stimulation with HLA matched CD8^+^ T cell epitopes (Panel d) or CMV gB overlapping pepmix (Panel e). Splenocytes were stimulated with HLA matched CD8^+^ CMV T cell epitopes or gB overlapping pepmix and then cultured for 10 days in the presence of IL-2. After incubation these T cells were assessed for IFN*γ* expression using intracellular cytokine assays. * P < 0.05; ** P < 0.01; *** P < 0.001 by Welch’s t test. Statistics are indicated in comparison with control group. If statistics are not indicated, then the comparison was not significant (ns).

**Fig 4:**
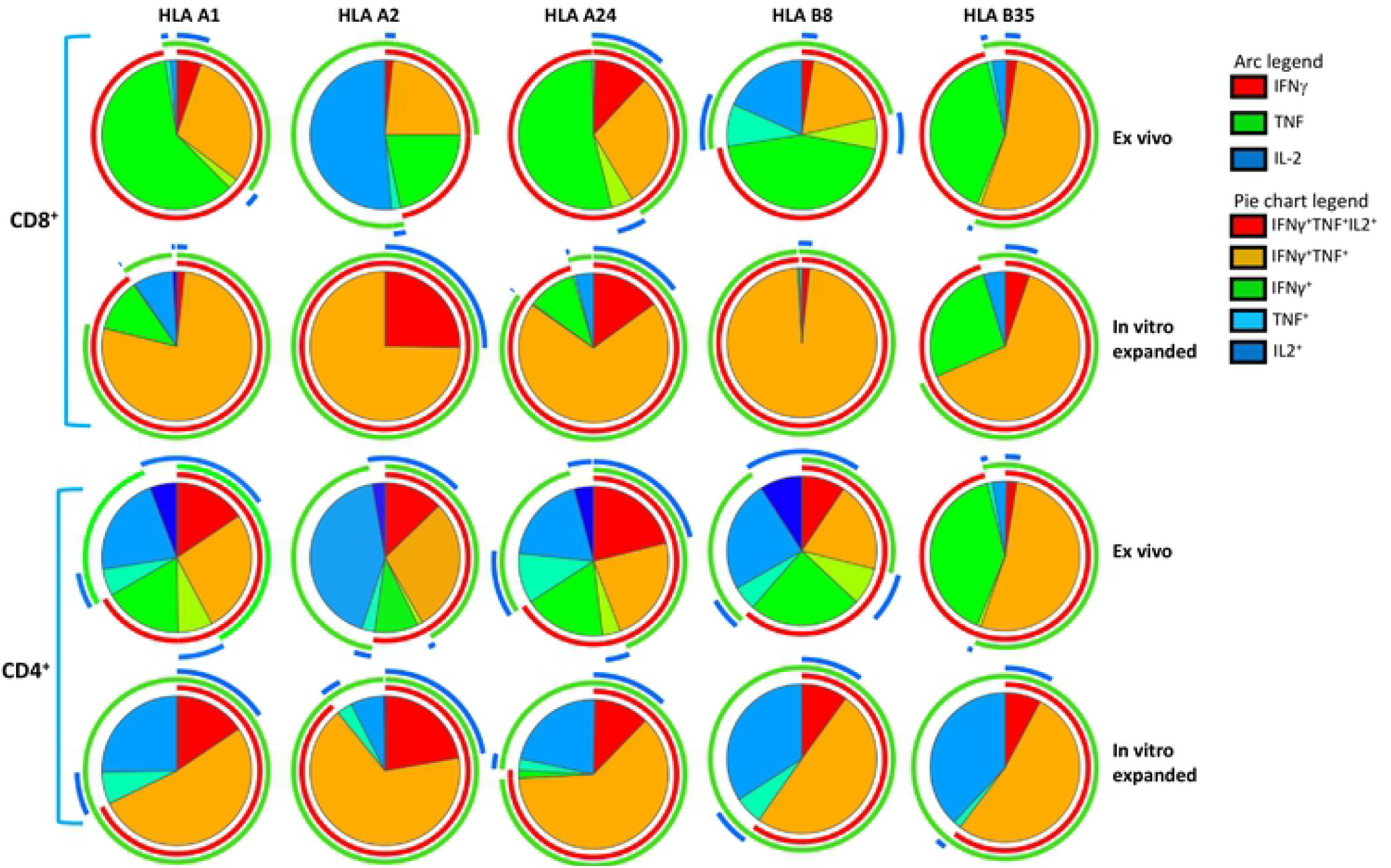
CMV vaccine induces polyfunctional CMV-specific CD8^+^ and CD4^+^ T cells: *In vitro* expanded CMV-specific T cells from HLA expressing immunized mice were assessed for polyfunctional profile using multiparametric intracellular cytokine assay. These T cells were assessed for the expression of IFN-γ, TNF and/or IL-2 simultaneously by flow cytometry in combination with SPICE software. Pie charts represent the proportion of CMV-specific CD8^+^ and CD4^+^ T cells expressing various combinations of IFNγ, TNF and/or IL2 cytokines after *ex vivo* and *in vitro* expansion. Arc indicates the proportion of CMV-specific CD8^+^ and CD4^+^ T cells expressing individual cytokines (IFNγ, TNF or IL-2).

To assess whether CMV vaccine formulation can also induce robust humoral immune responses we evaluated serum anti-gB antibody titres using ELISA. Data presented in Fig. 5a shows that the CMV vaccine formulation induced high gB-specific antibody titres in all HLA expressing mice. Higher magnitude of gB-specific antibody responses were observed in HLA A1, A2 and A24 mice while HLA B8 and B35 mice showed slightly lower antibody titres. Antibody isotyping analysis of gB-specific humoral immune response showed broad Th1 like IgG2b subtype in all HLA expressing mice, IgG3 in HLA A24, B8, B35 and A1 mice, and IgG2a predominantly in HLA A2 and A24 mice (Fig 5b). In addition, strong T_H_2-like IgM and IgG1 also observed in all expressing mice with the exception of IgG1 in HLA A2 expressing mice (Fig. 5b). More importantly, these antibodies neutralised CMV and blocked viral infection in fibroblasts and epithelial cells (Fig. 5c & d). Collectively, comprehensive evaluation of immunogenicity of CMV vaccine formulation in multiple human HLA expressing mice demonstrates that the CMV vaccine formulation is capable of inducing robust CMV-specific CD4^+^ and CD8^+^ T cells responses and neutralising antibody responses.

**Fig 5:**
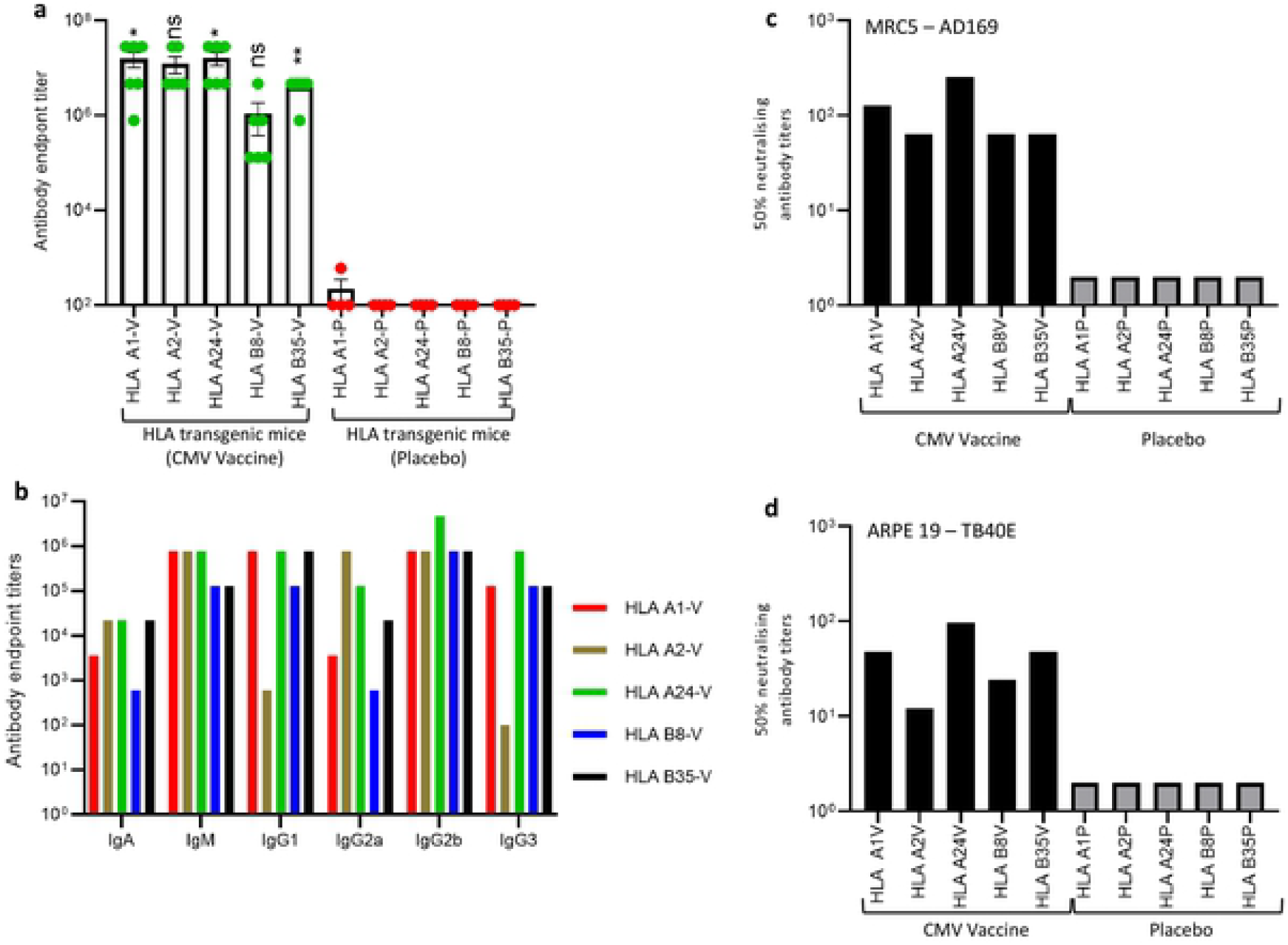
CMV vaccine induces gB-specific humoral immune responses in multiple human HLA expressing mice: HLA expressing mice (HLA A1, HLA 2, HLA A24, HLA B8 and HLA B35) were immunized with CMV vaccine (N=6) or placebo (N=4) as outlined in Fig 3a. Peripheral blood was collected on day 28 and assessed for gB-specific humoral immune responses using ELISA and CMV neutralisation assays. (a) CMV gB-specific Ig endpoint titres on day 28 in serum samples of HLA expressing mice immunized with CMV vaccine. (b) Prevalence of CMV gB-specific antibody isotypes (IgM, IgA, IgG1, IgG2a, IgG2b and IgG3) in pooled serum samples on day 28. (c and d) Neutralizing antibody titres in mice immunized with CMV vaccine. To determine the neutralising antibody responses, pooled serum samples from individual groups were serially diluted, and pre-incubated with CMV AD169 or TB40/E strains. MRC-5 or ARPE 19 cells were infected with serum-treated virus and virus infectivity was determined using an IE-1/IE-2 micro-neutralization assay. Error bars represent the mean ± SEM. *P < 0.05; **P < 0.01 by Welch’s t test. Statistics are indicated in comparison with control group. If statistics are not indicated, then the comparison was not significant (ns).

### Long-term CMV vaccine immunogenicity evaluation in HLA A24 expressing mice

One of the major challenges with previous CMV vaccines was the induction of long-term durable immunity and modest efficacy (22-24). In fact, durable vaccine-induced immune responses play a central role in minimising waning efficacy of vaccines against diverse pathogens. Therefore, it was important to test the long-term durability of CMV formulation induced immune responses and number of booster doses required to maintain persistent immune responses against CMV.

#### (a) Cellular Immune responses

In the next set of experiments, HLA A24 expressing mice were immunised with CMV vaccine or CpG 1018 adjuvant alone (placebo) formulations subcutaneously as outlined in Fig. 6a. These animals were systematically monitored for CMV-specific cellular and humoral immune responses on days 28, 42, 48, 84, 133, 203 and 217. *Ex vivo* analysis of splenocytes with HLA A24 restricted CD8^+^ T cell epitopes (AYA and QYD) showed that the CMV vaccine formulation induced significantly higher IFN-γ producing CMV-specific CD8^+^ T cell responses compared to the control and these robust T cell responses are maintained during the entire course of follow up (from day 28 to day 217). While IFN-γ producing CMV-specific CD8^+^ T cell responses were lower on day 42, there was a significant increase in antigen-specific CD8^+^ T cells with subsequent booster vaccination on days 42 and 210 (Fig. 6b).

**Fig 6:**
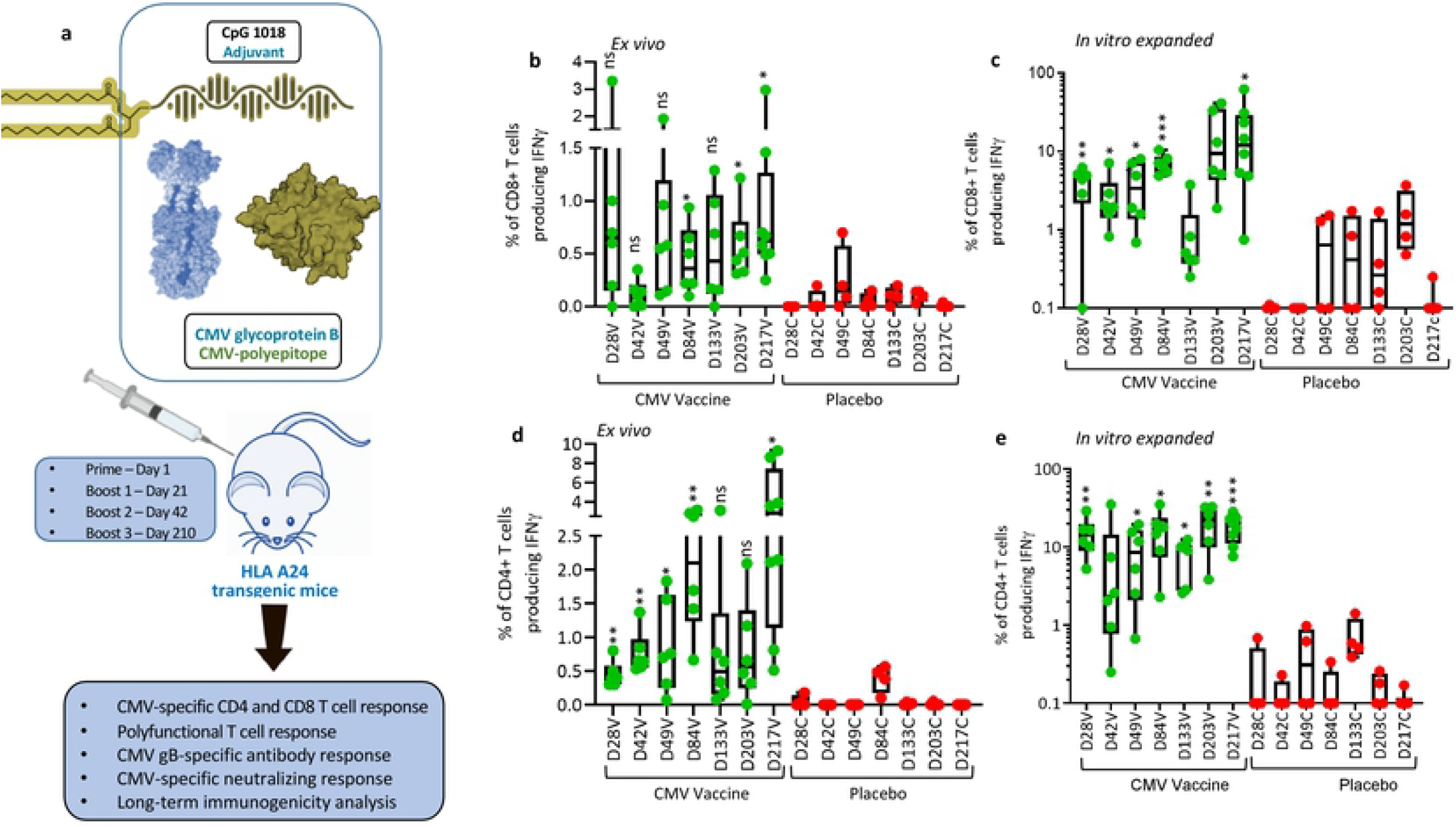
CMV vaccine induces long-term durable CMV-specific immunity: (a) Schematic outline of experimental plan to evaluate long-term durability of immune responses induced by CMV vaccine. HLA A24 expressing mice were immunized on day 0 and then boosted on days 21, 42 and 210 with CMV vaccine. Mice were sacrificed on days 28, 42, 49, 84, 133, 203 and 217 to perform longitudinal analysis of CMV-specific cellular and humoral immunity. (b and d) *Ex vivo* analysis of CMV-specific CD8^+^ and CD4^+^ T cell responses in HLA A24 expressing mice. Splenocytes from immunized and control mice were stimulated with HLA A24-restricted CD8^+^ T cell peptide epitopes (QYD & AYA) or with CMV gB overlapping peptides in the presence of brefeldin A and then assayed for IFNγ expression using intracellular cytokine assay. (c and e) Frequency of CMV-specific CD8+ and CD4+ T cells following *in vitro* expansion of antigen-specific T cells. Splenocytes from immunized mice were stimulated with HLA A24-restricted CD8^+^ T cell peptide epitopes (QYD & AYA) or with CMV gB overlapping peptides and cultured for 10 days in the presence of IL-2. On day 10, T cell specificity was assessed using intracellular cytokine assay. *P < 0.05; **P < 0.01; ***P < 0.001 by Welch’s t test. Statistics are indicated in comparison with control group. If statistics are not indicated, then the comparison was not significant (ns).

To confirm *ex vivo* cellular immune responses, splenocytes from vaccinated mice were stimulated with HLA A24-restricted peptide epitopes (AYA and QYD) and cultured for 10 days in the presence of recombinant IL-2 to allow expansion of antigen-specific T cells. We observed a significant expansion of antigen-specific CD8^+^ T cells with a mean frequency ranging from 1.9% and 20.05%, demonstrating the CMV vaccine formulation has potential to induce durable, long-term CD8^+^ T cell responses against CMV (Fig. 6c). In addition, the CMV vaccine formulation also induced high frequencies of IFN-γ producing CMV gB-specific CD4^+^ T cells compared to CpG 1018 alone with mean frequency ranging from 0.45% and 3.83% (Fig. 6d). There was a substantial increase in *ex vivo* IFN-γ producing CMV gB-specific CD4^+^ T cell with each booster vaccination on days 28, 49 and 84 (Fig. 6d). While these T cell responses slightly declined and plateaued on day 133 and 203, nevertheless after the third booster there was a robust increase in IFN-γ producing CMV gB-specific CD4^+^ T cell responses, indicating that long-term memory gB-specific CD4^+^ T cells can rapidly mature into effector cells after antigen reencounter (Fig. 6d). Additionally, *in vitro* stimulation of splenocytes with gB overlapping peptides induced expansion of IFN-γ producing CMV gB-specific CD4^+^ T cell responses at all the time points with a mean frequency ranging from 7.17% and 26.55% (Fig. 6e). Furthermore, longitudinal polyfunctional analysis revealed that at all the time points CMV-specific CD8^+^ and CD4^+^ T cells expressed multiple cytokines (Fig. 7). Majority of these antigen-specific T cells expressed IFNγ, TNF and IL-2 or IFN-γ and TNF. Taken together, these results demonstrate that CMV vaccine formulation based on CMVpoly and gB adjuvanted with CpG1018 was highly effective in inducing durable CMV-specific CD4^+^ and CD8^+^ T cell responses and majority of these effector cells displayed a polyfunctional profile.

**Fig 7:**
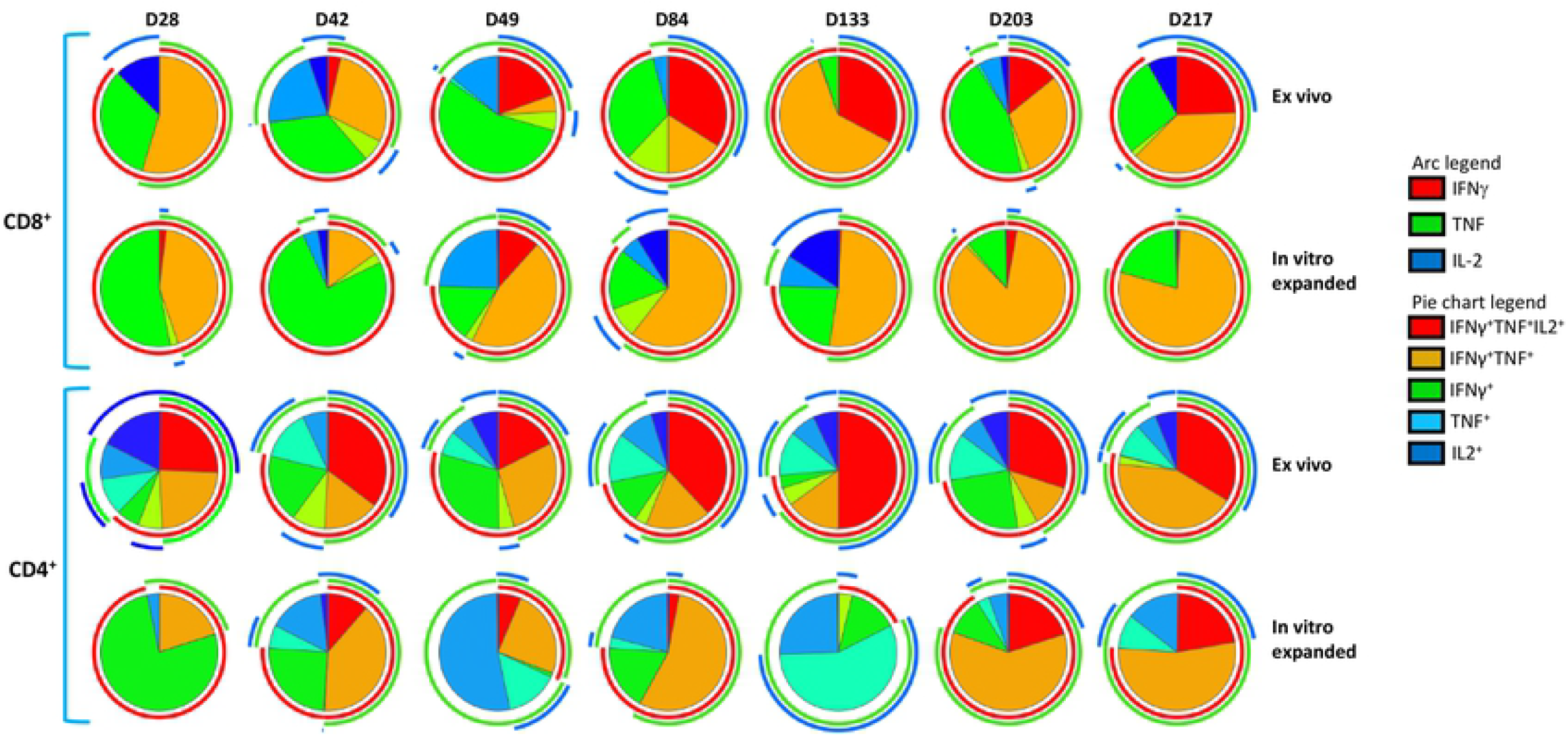
CMV vaccine induces long-term durable polyfunctional CMV-specific CD8+ and CD4+ T cell responses. *In vitro* expanded CMV-specific T cells from HLA A24 expressing mice were assessed for polyfunctional profile using multiparametric intracellular cytokine assay. These T cells were assessed for the expression of IFN-γ, TNF and/or IL-2 simultaneously by flow cytometry in combination with SPICE software. Pie charts represent the proportion of CMV-specific CD8^+^ and CD4^+^ T cells expressing various combinations of IFNγ, TNF and/or IL2 cytokines after *ex vivo* and *in vitro* expansion. Arc indicates the proportion of CMV-specific CD8^+^ and CD4^+^ T cells expressing individual cytokines (IFNγ, TNF or IL-2).

#### (b) Humoral immune responses

Germinal centres plays a vital role in induction of memory B cells and long-lived plasma cell to secrete the high-affinity antibodies required for long-term serological immunity (25). Data presented in Fig 8a & b shows that the CMV vaccine induced a significantly higher proportion of germinal centre B cells (B220^+^GL7^+^Fas^+^) on days 28, 49 and 84 with a mean frequency ranging from 3.15% to 4.07% and then declined thereafter. Assessment of CMV gB-specific IgG secreting plasma cells in spleens by ELISpot assay indicated that the CMV vaccine formulation induced a significantly higher and sustainable plasma B cell response at all time points with mean frequencies ranging from 73.3 to 357.7 antibody secreting cells/3×10^5^ splenocytes (Fig. 8c). The CMV vaccine also induced long-term gB-specific memory B cell responses with mean frequency ranging from 677 and 6916 antibody secreting cells/3×10^5^ splenocytes (Fig. 8d). Interestingly, a dramatic increase in memory B cells was observed following third booster vaccination (Day 217) (Fig. 8c & d). These observations were consistent with detection of high levels of antibody response against gB protein (Fig. 9a). Characterisation of antibody responses revealed that CMV vaccine formulation induced durable and broad T_H_1 isotypes (IgG2a, IgG2b and IgG3) (Fig 9b-g). More importantly, these antibodies neutralised CMV and blocked viral entry in fibroblasts and epithelial cells (Fig. 10a & b). Furthermore, serum antibodies from mice immunised with CMV vaccine formulation exhibited strong binding to cell-associated gB on CMV-infected cells (Fi. 10c & d). This antibody-mediated binding was significantly higher immediately following booster vaccination on days 28, 49 and 217. Finally, emerging data shows that the gB protein used in gB/MF59 vaccine was in monomeric form, but in naturally infected individuals the gB assumes its native trimeric conformation. Thus, to determine the immunogenicity of the gB multimer we have performed Western blot analysis using mouse serum. Interestingly, data obtained from Western blot analysis indicated that sera from mice immunised with CMV vaccine weakly reacted to gB monomer; however, a strong reaction was observed to gB trimer, indicating that gB trimer is more immunogenic (Fig. 10e). Collectively, these data indicate that CMV vaccine formulation demonstrated its potential to induce durable and qualitative CMV-specific humoral and cellular immune responses.

**Fig. 8:**
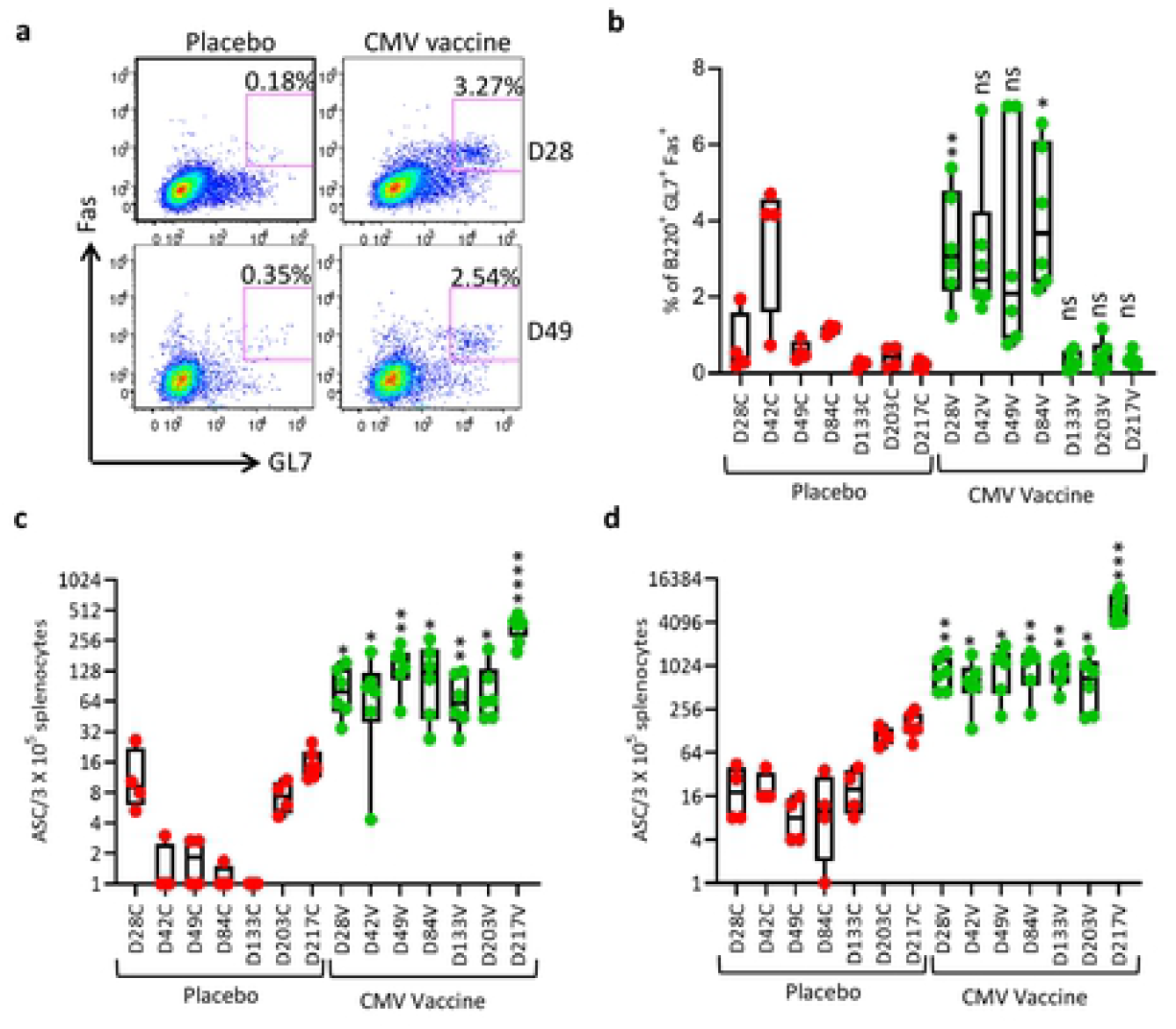
CMV vaccine induces long-term durable GC B cell and CMV gB-specific antibody secreting B cell responses: HLA A24 expressing mice immunized with CMV vaccine as outlined in Fig. 6a and assessed for GC B cells and CMV gB-specific antibody secreting B cell responses. To determine the GC B cell responses, single cell suspensions were prepared from spleens and then stained with PE conjugated anti-B220, FITC conjugated anti-GL7 and APC conjugated anti-CD95 antibodies. (a) Representative FACS plots of B220^+^GL7^+^Fas^+^ B cells in spleens of mice immunised with CMV vaccine or placebo on day 28 and 49. (b) Frequencies of B220^+^GL7^+^Fas^+^ B cells in spleens of mice immunised with CMV vaccine or placebo from day 28 to 217. (c and d) *Ex vivo* and memory antibody secreting B cells in splenocytes of mice immunised with CMV vaccine or placebo on days 28, 42, 49, 84, 133, 203 and 217 using ELISpot assay. *P < 0.05; **P < 0.01; ***P < 0.001, ****P < 0.0001 by Welch’s t test. Statistics are indicated in comparison with control group. If statistics are not indicated, then the comparison was not significant (ns).

**Fig 9:**
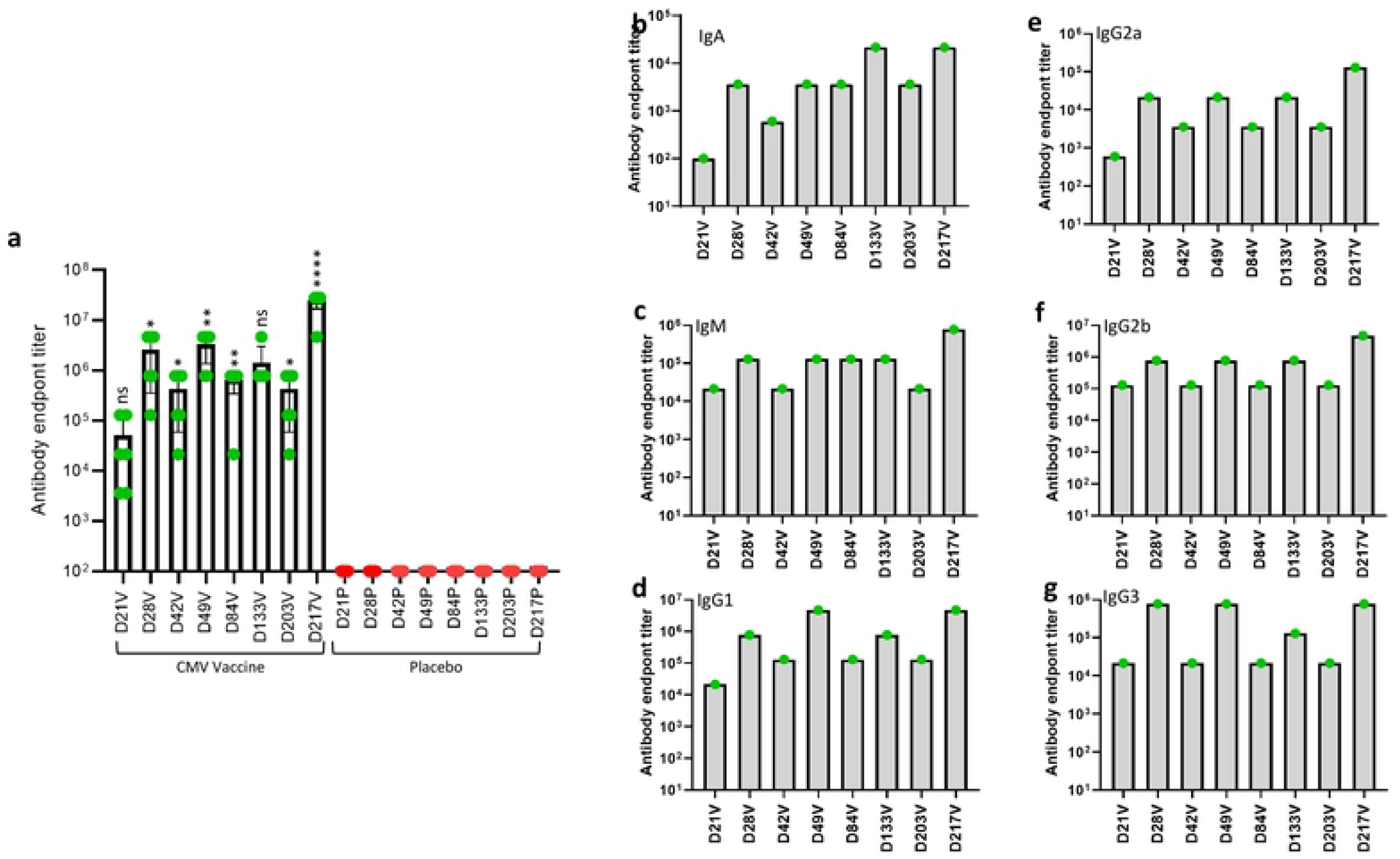
CMV vaccine formulation induces sustainable CMV gB-specific antibody responses: HLA A24 expressing mice were immunized with CMV vaccine or placebo as outlined in Fig. 6a. Serum samples were collected from these animals and analysed for gB-specific antibody titres using ELISA. (a) CMV gB-specific Ig endpoint titres in serum samples of mice immunised with CMV vaccine or placebo on day 21, 28, 42, 49, 84, 133, 203 and 217. (b - g) shows prevalence of CMV-gB specific immunoglobulin isotypes (in pooled serum samples) (IgA, IgM, IgG1, IgG2a, IgG2b and IgG3) induced following immunisation with CMV vaccine formulation or placebo on day 21, 28, 42, 49, 84, 133, 203 and 217. *P < 0.05; **P < 0.01; ***P < 0.001, ****P < 0.0001 by Welch’s t test. Statistics are indicated in comparison with control group. If statistics are not indicated, then the comparison was not significant (ns).

**Fig 10:**
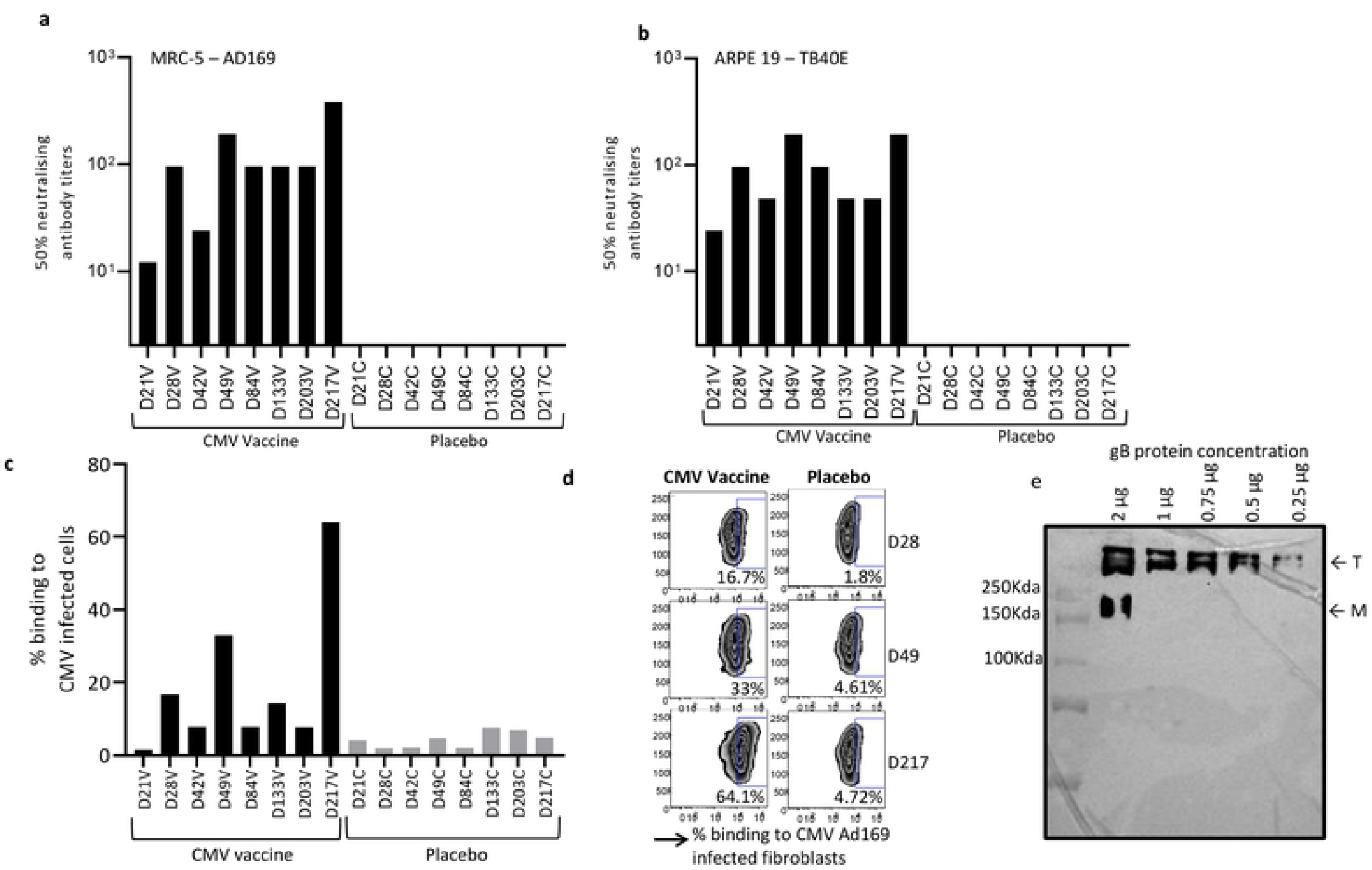
Functional characterisation of CMV vaccine formulation induced antibody responses: HLA A24 expressing mice were immunized with CMV vaccine as outlined in Fig 6a. Serum samples were collected at each time point and assessed for CMV neutralising antibody responses using microneutralisation assay. Neutralising antibody responses were measured by IE-1 staining in fibroblasts infected with CMV AD 169 and in ARPE 19 cells infected with CMV TB40/E strain. (a and b) Neutralising antibody titres induced following immunisation with CMV vaccine or placebo on day 21, 28, 42, 49, 84, 133, 203 and 217 against CMV AD 169 and TB40/E strains. (c and d) Flow cytometry analysis of binding of gB-specific antibodies to fibroblasts infected with CMV AD169 strain. (e) Western blot analysis of antibodies induced following immunisation with CMV vaccine or placebo binding to CMV gB monomer (M) or trimer (T).

## Discussion

CMV is medically a significant human pathogen and vaccination to reduce congenital infection and CMV-associated disease in transplant patients has been proposed as a potential medical intervention. Our goal in this study was to develop a novel protein based subunit vaccine to induce strong and durable CMV-specific humoral and cellular immune responses. Historically, the induction of potent neutralising antibody responses have been considered as a benchmark for evaluation of vaccine efficacy; however, CMV is not amenable to antibody-dependent immunity alone (26-28). Cellular immune responses are also needed as CMV-specific CD4^+^ and CD8^+^ T cells play an important role in reducing virus acquisition, prolonged virus shedding, viral replication and CMV-associated disease in congenital and transplant patients (11, 14, 20, 29). Therefore we hypothesised that a vaccine designed to induce both humoral and cellular immune responses against multiple CMV antigens may offer better protection. To target multiple immunodominant antigens, we have engineered a novel, artificial CMV-specific antigen; CMVpoly protein. CMVpoly is designed to encode 20 different CD8^+^ T cell epitopes derived from five different CMV immunodominant antigens which are expressed in various stages of virus replication, therefore it induces a robust and broad repertoire of CMV-specific CD8^+^ T cell responses. *In vitro* immunogenicity analysis indicated that CMVpoly triggered robust expansion of polyfunctional CMV-specific CD8^+^ T cells targeting multiple antigens from seropositive healthy donor PBMC and expanded cells were able to recognise and kill CMV infected cells. Antiviral activity of CMV-specific CD8^+^ T cells is considered as an important aspect in CMV vaccine development because these effector cells can eliminate virus infected cells and restrict CMV spread via direct cell-to-cell contact (30).

Another focus of this study was to improve immunogenicity of gB protein because CMV gB has been considered as an essential component of prophylactic vaccines due to its crucial role in early stages of viral infection, the cell-to-cell spread of intracellular virions and the fusion of infected cells leading to formation of multinucleated cells (31). In this study, we have successfully expressed gB protein consisting of extracellular and intracellular domains in CHO cells and purified using a combination of chromatography techniques. The biophysical characterisation of purified gB protein revealed that CMV gB predominantly existed as a trimer or dimer of a trimer in solution. During natural infection native gB homotrimer is required for the fusion activity; therefore, for induction of neutralising antibody responses gB in its trimeric form is considered as the ideal conformation in vaccine development (32, 33). In addition, emerging data shows that trimeric gB induces higher serum titres of gB-specific IgG antibody and neutralising antibody responses compared to monomeric CMV gB (34).

We combined CMVpoly and oligomeric gB with human compatible adjuvant CpG-ODN 1018 and assessed its immunogenicity in multiple HLA expressing mice. Our initial studies demonstrated that the two doses of CMV vaccine induced strong ex vivo and memory multi-antigen-specific CD8^+^ and gB-specific CD4^+^ T cell responses and neutralising antibody responses against AD169 and TB40E CMV strains. More importantly, the CMV-specific T cells from vaccinated mice displayed polyfunctional profiles secreting three (IFNγ^+^TNF^+^IL2^+^) or two (IFNγ^+^TNF^+^) cytokines. Previously published studies have shown that protective immunity of CD4^+^ and CD8^+^ T cells is associated with production of multiple effector cytokines and impaired polyfunctionality of CMV-specific T cells can compromise control of CMV replication in children with congenital CMV infection (19-21).

One of the important attributes for an effective CMV vaccine is generation of durable memory immune responses as protection against CMV in congenital and transplant settings is needed throughout women of child bearing age or course of transplantation. Pre-existing antibody responses are usually considered as a primary line of defence and crucial for prevention of primary infection or reinfection at mucosal surfaces. CMV is a highly cell-associated virus; thus, cytotoxic T cell responses are also important for mediating suppression of viral replication and reactivation by elimination of virus infected cells (35). Thus, the magnitude of immune response generated through natural immunity to CMV remains an important benchmark for evaluation of efficacy of CMV vaccine. Our long-term follow up studies showed that the CMV vaccine induced durable antigen-specific CD8^+^ T cell, gB-specific CD4^+^ T cell, B cell and neutralising antibody responses which were maintained beyond day 210. Interestingly, a clear enhancement of both cellular and humoral immunity was observed after booster immunisations on days 28, 49 and 210. Furthermore, we have also shown that gB-specific antibodies strongly bound to both soluble trimeric gB and gB protein expressed on CMV infected cells. These observations are consistent with recent finding that the protective ability of gB-specific antibodies binding to cell-associated gB correlated with gB/MF59 vaccine efficacy (36).

Collectively, the bivalent CMV vaccine formulated with CMVpoly and gB proteins with CpG1018 adjuvant is highly immunogenic in multiple HLA expressing mice and can induce durable and multi-effector humoral and cellular immune responses against CMV. These results clearly support further development of this bivalent subunit CMV vaccine to test safety, immunogenicity and efficacy in humans.

## Materials and Methods

### CMVpoly20PL-NH protein expression and purification

Chemically competent *E. coli* BL21-codonPlus (DE3) RP cells (Agilent Technologies) were transformed with CMVpoly expression vector (pJ404-Atum Bio). Transformed cells were plated on Luria Bertani (LB) agar supplemented with ampicillin (LB-Amp) 100 μg/mL and plates were incubated overnight at 37°C. An isolated colony was picked and inoculated into 10 ml of terrific broth containing 100 μg/mL ampicillin (TB-Amp broth) and grown in a shaker with 37° C and 200 rpm for overnight. A small amount of overnight grown culture was inoculated into 50 mL of TB-Amp broth and grown for 12 hours. About 1% of culture from 50 mL culture transferred into 3 L of TB-Amp broth and culture was grown until OD reached to 0.6 at 600nm. CMVpoly20PL-NH induction was carried out by adding 1 mM/mL of IPTG. These cells were allowed to grow for an additional 4 hours and protein expression levels were determined by analysing un-induced and induced samples on the 12% sodium dodecyl sulphate polyacrylamide gel electrophoresis (SDS-PAGE).

At the end of the induction phase, *E. coli* culture was harvested by centrifugation at 13,000 rpm for 15 minutes, cell pellet was resuspended in cell lysis buffer (25 mM Tris pH 7.5, 5 mM EDTA, 0.5% TritonX 100, 0.5 mg/mL lysozyme) supplemented with a protease inhibitor cocktail (Roche, Mannheim, Germany) and cells were lysed using sonicator The lysate was analysed on SDS-PAGE and then clarified by centrifugation. Following SDS-PAGE analysis protein was found to be in the pellet in the form of inclusion bodies (IBs). To eliminate the host protein contaminants IBs were washed with TE buffer (25mM Tris and 5mM EDTA pH 7.5) and then with 100 mM NaH_2_PO4, 10 mM Tris, 4 M urea pH 7.5 buffer. Purified IBs were solubilised in in of 100 mM NaH2PO4, 10 mM Tris, 2.5 mM DTT, 8 M urea pH 5.5 buffer, the soluble protein was clarified by centrifugation CMVpoly protein was purified using SP-Sepharose and Q Sepharose matrix. Purified protein was dialysed against 25 mM glycine buffer pH 3.8, protein was concentrated and filtered through Mustang E membrane (PALL Corporation, NY, USA) to further remove endotoxin contaminants, protein was filter sterilised using 0.22μ membrane filter and then total protein was estimated using BIO-RAD Bradford protein assay kit and UV absorbance at 280 nm and purified protein was stored at -70°C.

### gB protein expression and purification

The coding sequence of HCMV gB expressing extracellular and intracellular domains from HCMV strain AD169, was codon optimised for mammalian expression to enhance the protein expression and cloned into a mammalian expression vector (Geneart). The native N-terminal signal sequence (i.e., amino acids 1 to 31 of the sequence) was replaced with the heterologous IgG heavy chain signal peptide to secrete the expressed polypeptide into cell culture supernatant. The encoded sequence of the furin cleavage site was mutated from Arg456 to Gln, Arg458 to Thr and Arg459 to Gln. Chinese Hamster Ovary (CHO) K1 cells were transfected with the modified gB polypeptide plasmid and stable cells expressing modified gB polypeptide were selected with 9 µg/mL blasticidin and 400 µg/mL Zeocin. Modified gB polypeptide was expressed in a 10 L bioreactor in fed-batch mode. On day 12, cell culture was harvested and centrifuged to separate supernatant from cell debris. The supernatant was buffer exchanged into 200 mM Tris-HCL (pH 8.0) and then loaded on Poros 50HQ resin (anion exchange chromatography). The column was washed with 20 mM Tris-HCL, 70 mM NaCl (pH 8.0) buffer, before eluting the protein with 20 mM Tris-HCL, 180 mM NaCl (pH 8.0) buffer. Eluted polypeptide was buffer exchanged with 5 mM phosphate buffer (pH 7.0) and passed through a ceramic hydroxyapatite (CHT) Type II to eliminate host cell protein contaminates. To further improve the purity, the modified gB polypeptide was buffer exchanged into 50 mM sodium acetate (NaOAc) (pH 5.0) buffer and then loaded on POROS XS (cation exchange chromatography) column. The modified gB polypeptide bound to the POROS XS column was eluted with 50 mM NaOAc, 500 mM NaCl (pH 5.0) buffer and then buffer exchanged against 25 mM glycine (pH 4.0) buffer. The final purified polypeptide concentration was determined by UV280 using extinction coefficient 1.209, analysed on SDS-PAGE gel and stored at -70<u>°</u>C.

### Mass photometry analysis of CMV gB protein

Mass photometry analysis was performed using a Refeyn OneMP instrument (Refeyn Ltd). 10 µL of buffer was applied and then 1 µL of gB sample at 0.04 mg/mL was applied and 5965 frames were recorded.

### Negative stain electron microscopy

The gB protein was diluted to 0.004 mg/mL and 5 µL applied to glow-discharged carbon-coated Formvar grids (ProSciTech, Australia), stained with 2% (w/v) uranyl acetate for two minutes, then blotted and air-dried for 10 minutes. Samples were imaged at 150 000x magnification using a JEOL JEM 1011 TEM.

### CMVpoly immunogenicity evaluation *in vitro*

To determine the immunogenicity of CMVpoly protein, 6 × 10^6^ PBMC from 10 different healthy CMV seropositive donors were stimulated with 20 µg of CMVpoly protein. Cells were cultured for 14 days in the presence of IL-2. To determine the antigen specificity expanded PBMCS were re-stimulated with the individual antigen peptides from CMVpoly in an IFNγ ICS assay.

### *In vitro* cytotoxicity assay

To determine the ability of CMV-specific CD8+ T cells to kill CMV infected fibroblasts, 6 × 10^6^ PBMC from 3 different healthy CMV seropositive donors were stimulated with 20 µg of CMVpoly protein. Cells were cultured for 14 days in the presence of IL2. HLA restriction of the donor 1 was HLA A 01, A3, B7 and B8; donor 2 HLA A1, A2 and B8 and donor 3 HLA A2, A24 and B40. Antigen specificity of *in vitro* expanded CMV-specific CD8+ T cells were determined using polyfunctional ICS assay. Fibroblasts expressing HLA A*01:01, A*02:01 and B*08:02 were plated and then treated with IFNγ for upregulation of HLA class I molecules. Fibroblasts were infected with CMV TB40/E strain overnight to allow CMV antigen expression. Uninfected fibroblasts also included as a control. Fibroblasts infected with CMV or uninfected fibroblasts were co-cultured with in vitro expanded CMV-specific CD8+ T cells at 1:1 effector to target ratio for 48 hours in Real Time Cell Analyser (RTCA), xCELLigence (Agilent Technologies Inc, Santa Clara, CA, USA). Cytolysis percentage was calculated using RTCA software.

### Evaluation of immunogenicity of CMV vaccine formulated with various combinations of adjuvants

All mouse studies were approved by QIMR Berghofer animal ethics committee. All human HLA expressing mice (HLA A1, HLA A2, HLA A24, HLA B8 and HLA B35) were bred and maintained under pathogen-free environment at the QIMR Berghofer. These expressing mice are deficient in expressing mouse MHC class I molecule and contain transgenes of the commonly expressed human HLA class I molecules. In order to evaluate the immunogenic response to the CMVpoly protein and CMV glycoprotein B (gB), proteins were admixed with unmethylated CpG oligodeoxynucleotides 1018 (CpG 1018) (TriLink BioTechnologies). Six to eight week old mice were immunized subcutaneously (s.c.) with 100 µL volume at the base of the tail with CMV vaccine formulation containing 30 µg of CMVpoly and 5 µg of gB protein admixed with 50 µg of CpG 1018 (CMV vaccine formulation). Mice immunised with 50 µg of CpG 1018 adjuvant alone was used as a negative controls (placebo). Mice were tail bled on day 21 and a booster dose was given. Mice were then sacrificed in day 28 and evaluated for immune responses.

To assess the long-term durability of CMV vaccine induced immune responses HLA A24 human HLA expressing mice were immunised with CMV vaccine formulation containing 30 µg of CMVpoly and 5 µg of gB protein admixed with 50 µg of CpG 1018 on day 0 and then boosted on day 21, 42 and 210. Control group mice were injected with CpG1018 adjuvant alone. Mice were sacrificed on day 28, 42, 49, 84, 133, 203 and 217 to evaluate long-term CMV-specific immune responses.

### Intracellular cytokine staining to assess IFN-γ, multiple cytokines, germinal centre B cells or T follicular helper cells responses

Following vaccination, splenocytes were stimulated with 0.2 μg/mL of HLA matching CMV CD8+ T cell peptides (HLA A1-VTE & YSE; HLA A2-NLV, VLE & YIL; HLA A24-QYD & AYA; HLA B8-QIK, ELR & ELK; HLA B35-FPT & IPS) to determine the CMV-specific CD8+ T cell response or with 0.2 μg/mL of gB pepmix (gB overlapping peptides-15mers with 11 aa overlap) to detect CMV-specific CD4+ T cell response in the presence of GolgiPlug^®^ and GolgiStop (BD PharMingen) for 6 hours, cells were washed twice, then incubated with APC-conjugated anti-CD3, FITC-conjugated anti-CD4 and PerCP conjugated anti-CD8. Cells were fixed and permeabilized using a BD Cytofix/Cytoperm kit, then incubated with PE conjugated anti-IFN-γ. To assess the expression of multiple cytokines, cells were stained with PerCP conjugated anti-CD8 and BV786 anti-CD4 surface markers and then intracellularly with PE-conjugated anti-IFN-γ, PE-Cy7 conjugated anti-TNF, and APC conjugated anti-IL2. To assess the germinal centre B cell response splenocytes were stained with PE conjugated anti-B220, FITC conjugated anti-GL7 and APC conjugated anti-CD95. Cells were acquired on a BD FACSCanto II and data was analysed using FlowJo software (Tree Star).

### *In vitro* expansion of CMV-specific CD4+ and CD8+ T cells following vaccination

Following vaccination, 5 × 10^6^ Splenocytes from immunized mice were isolated and stimulated with 0.2 μg/mL of matching CMV CD8+ T cell peptides or gB pepmix (gB overlapping peptides-15mers with 11 aa overlap) and cell were cultured in a 24 well plate for 10 days at 37 °C 10% CO_2._ Cultures were supplemented with recombinant IL-2 on days 3 and 6 and on day 10 and T cell specificity was assessed using ICS assay.

### Mouse IgG ELISpot assay

To measure ex vivo gB-specific antibody secreting cells, PVDF ELISpot plates (Millipore) were treated with 70% ethanol. Plates were washed five times with distilled water, coated with 100 µL/well CMV gB protein (25 µg/mL) or anti-IgG antibody (15 µg/mL) and incubated overnight at 4° C. Plates were blocked with DMEM containing 10% serum, 300,000 cells/well in triplicates from each mouse was added and then incubated for 18 hours in a 37° C humidified incubator with 5% CO_2_. Cells were removed and plates were washed. Detection antibody anti-IgG conjugated to HRP (MABTECH) was added and incubated for 2 hours at room temperature. Plates were washed; Streptavidin-ALP was added and incubated at room temperature for 1 hour followed by washing and treating plates with substrate solution containing BCIP/NBT (Sigma-Aldrich) until colour development is prominent. Colour development was stopped by washing plates with water and plates were kept for drying overnight. To measure memory B cell response, the spleen cells (5 × 10^5^) were activated with a mixture of R484 and recombinant mouse IL-2 for five days in 24 well plate and then ELISpot was carried out as stated above. Number of spots were counted in an ELISpot reader.

### ELISA

Serum total anti-gB antibody and antibody isotype titres were evaluated by an enzyme-linked immunosorbent assay (ELISA). Briefly, 96-well plates pre-coated with 50µL of recombinant HCMV gB protein (2.5 µg/mL of gB protein diluted in phosphate buffer saline) and plates were incubated at 4° C overnight. Plates were washed with phosphate buffer saline containing 0.05% Tween 20 (PBST) buffer and then blocked with 5% skim milk. The serially diluted serum samples (day 21 or day 28) were added and incubated for 2 hours at room temperature. After washing with PBST, plates were incubated with HRP-conjugated sheep anti-mouse Ig antibody (to determine total antibody response) or HRP-conjugated goat anti-mouse IgA, IgM, IgG1, IgG2a, IgG2b or IgG3 antibody (SouthernBiotech) (to determine antibody isotype) for 1 hour. These plates were washed and incubated with 3,3’,5,5’-Tetramethylbenzidine substrate solution for 10 mins and then colour development was stopped by adding 1 N HCl. OD at 450 nm was measured using an ELISA reader.

### Microneutralisation assay

Neutralizing activity was determined against AD169 and TB40/E strains of CMV. Human fibroblast MRC-5 or human adult retinal pigment epithelial cells (ARPE-19) were plated in 96-well flat-bottomed plates. The next day, serum samples from mice vaccinated with CMV vaccine formulation were serially diluted and added to a standard number of virus particles (1000 p.f.u. per well) diluted in DMEM with no serum in 96-well U-bottomed plates and incubated for 2 h at 37° C and 5% CO_2_. As a positive control, virus without serum and a negative-control serum without virus were also included in the test. The serum/CMV mixture was then added to the MRC5 and ARPE-19 cells and incubated at 37° C and 5% CO2 for 2 hours. After incubation, the mixture was discarded and the cells washed gently five times with DMEM containing 10% FCS (D10) and a final volume of 200 ml R10 was added to each well, followed by incubation for 16–18 h at 37° C and 5% CO_2_. The cells were fixed with 100 ml chilled methanol and incubated with Peroxidase Block (Dako) followed by mouse anti-CMV IE-1/IE2 mAb (Chemicon) at room temperature for 3 hours. Cells were then incubated with 50 ml HRP-conjugated goat anti-mouse Ig (diluted 1:200 in PBS) per well for 3 hours at room temperature. The cells were stained with 20 ml diaminobenzidine plus substrate (Dako) per well for 10 min at room temperature and positive nuclei that stained dark brown were counted. The neutralizing titre was calculated as the reciprocal of the serum dilution that gave 50% inhibition of IE-1/IE-2-expressing nuclei.

### Western blot analysis

To determine the most immunogenic form of oligomeric gB porotein a Western blot analysis was performed. The oligomeric forms of purified gB purified protein conentrations ranging from 2 to 0.25 µg were separated on 8% SDS-PAGE under non-reducing conditions. Following gB protein resolution on SDS-PAGE, the gB protein was transferred to Hybond-C nitrocellulose membrane. After transfer, membranes were washed, blocked and the probed withof mouse serum (1:3000 dilution). Membranes were washed and then incubated with HRP-conjugated goat anti-mouse Ig antibody, followed by a wash and incubation with Immobilon ECL ultra western HRP substrate. The signal was captured using Invitrogen CL1500 Chemi Gel Doc System.

### CMV gB-specific antibodies binding to cell-associated gB on CMV-infected fibroblasts assay

Human fibroblasts cell line, Mrc-5 cells were grown to 50% confluency in a T75 flask. Cell were infected with CMV AD169 strain at multiplicity of infection (MOI) of 2.0 at 37° C and 5% CO_2_ for 2 hours. Following infection cells were washed and incubated with DMEM containing 10% FCS for 48 hours to allow cell-cell virus spared. Infected cells were washed with PBS and then cells were dislodged with trypsin-EDTA. Cells were washed, counted and then resuspended at 10^6^ viable cells/mL. Cells were stained with cell trace violet and incubated for 20 minutes at room temperature. Cells were washed and then fixed with 4% paraformaldehyde for 10 minutes at room temperature. Cells were washed twice, plated 20,000/well in 96-well V-bottom plates, cells were pelleted by centrifugation and supernatant was discarded. Mouse serum samples obtained from HLA A24 human expressing mice following immunization with CMV vaccine or place on day 49 were polled and then diluted to 1:512. Diluted serum samples were added to the Mrc-5 cells and incubated for 2 hours at 37<u>°</u>C and 5% CO_2_. Cells were washed and then stained with anti-mouse AF488 IgG (H+L) for 30 minutes at 4°C. Cells were acquired on a BD FACSCanto II and data was analyzed using FlowJo software (Tree Star). The percentage of CMV gB-specific antibody binding to CMV infected fibroblasts was calculated from the percentage of viable AF 488 positive cells.

## Acknowledgements

The authors acknowledge the facilities, and the scientific and technical assistance, of the Microscopy Australia Facility at the Centre for Microscopy and Microanalysis (CMM), The University of Queensland.

